# Serotonin neurons in mating female mice are activated by male ejaculation

**DOI:** 10.1101/2023.05.14.540716

**Authors:** Eileen L. Troconis, Changwoo Seo, Akash Guru, Melissa R. Warden

## Abstract

Sexual stimulation triggers changes in female physiology and behavior, including sexual satiety and preparing the uterus for pregnancy. Serotonin is an important regulator of reproductive physiology and sexual receptivity, but the relationship between sexual stimulation and serotonin neural activity in females is poorly understood. Here, we investigated dorsal raphe serotonin neural activity in females during sexual behavior. We found that serotonin neural activity in mating females peaked specifically upon male ejaculation, and remained elevated above baseline until disengagement. Artificial intravaginal mechanical stimulation was sufficient to elicit increased 5-HT neural activity but the delivery of ejaculatory fluids was not. Distal penis erectile enlargement (“penile cupping”) at ejaculation and forceful expulsion of ejaculatory fluid each provided sufficient mechanical stimulation to elicit serotonin neuron activation. Our study identifies a female ejaculation-specific signal in a major neuromodulatory system and shows that intravaginal mechanosensory stimulation is necessary and sufficient to drive this signal.

## INTRODUCTION

For species with sexual reproduction and internal fertilization, mating facilitates the transfer of sperm into the female body. During the appetitive and consummatory phases of mating, the female receives multimodal sexual stimulation that is independent of sperm transfer but just as critical for successful reproduction (Hashikawa et al., 2016; Lorenz, 1981; Lenschow and Lima, 2020; Wei et al., 2021; Yin and Lin, 2023). Visual, acoustic, and olfactory stimuli in the appetitive phase are necessary for conspecific recognition, evaluation of mate quality, and courtship (Coen et al., 2014; Egnor and Seagraves, 2016; Ishii et al., 2017; Osakada et al., 2018; Ribeiro et al., 2018). Here we focus on the consummatory phase, in which mechanosensory and chemosensory signals provided by the male elicit dramatic changes in female physiology and behavior.

The rodent copulatory sequence is characterized by a number of temporally spaced mounts and intromissions with an ejaculation during the final intromission, a sequence that can be repeated several times (Beach, 1956; McClintock and Adler, 1978; McClintock et al., 1982). There is a decline in sexual receptivity immediately following copulation, and more intense vaginocervical stimulation leads to a lengthier period of reduced receptivity. The time it takes a female rat to return to the male between mounts rises with the number of intromissions she receives, and a female that has received intromission with ejaculation takes longer to return to the male than a female that has received only intromission (Peirce and Nuttall, 1961; Bermant, 1961; Bermant and Westbrook, 1966; Hill and Thomas, 1973; Erskine, 1985; Yang and Clemens, 1998). Mechanical vaginocervical stimulation also elicits behavioral changes such as lordosis and immobility during mounting (Komisaruk and Diakow, 1973; Naggar and Komisaruk, 1977; Lehmann and Erskine, 2004).

Mechanical vaginocervical stimulation is also essential for inducing physiological changes that initiate and support early pregnancy and pseudopregnancy in rodents. In rats, repeated intromission without ejaculation is sufficient to induce pseudopregnancy, but in mice, ejaculation is required (McGill and Coughlin, 1970). If mating mice are separated just before penile cupping and fluid expulsion most females do not become pseudopregnant, but if separation happens a few seconds later most females do become pseudopregnant. Ejaculatory fluid expulsion and penile cupping provide enough intravaginal mechanical stimulation to induce pseudopregnancy in mice, even without the formation of the copulatory plug (McGill et al., 1968). The expulsion of the ejaculatory fluid has been characterized as “forceful” (McGill and Coughlin, 1970)—the pressure inside the mouse penis corpus spongiosum at ejaculation is more than twice the pressure during pre-ejaculatory intromissions (Soukhova-O’Hare et al., 2007). In humans the pressure exerted by the ischiocavernosus and bulbospongiosus muscles is sufficient to expel the ejaculate 30 to 60 cm (Masters and Johnson, 1966). The neural mechanisms that underlie the translation of these brief but intense mechanosensory stimuli into persistent behavioral and hormonal state changes are currently unclear.

In addition to mechanosensory stimulation, chemical signaling through the ejaculatory fluid is a common mechanism by which female post-copulatory reproductive function is regulated across species. For example, after mating *D. melanogaster* females become sexually unreceptive and begin laying eggs (Auer and Benton, 2016), a critical behavioral switch that is primarily triggered by the sex peptide from the male (Wang et al., 2020; Yapici et al., 2008). In addition, seminal fluid in *D. melanogaster* (Heifetz and Wolfner, 2004) contains signaling molecules that condition the environment in the female reproductive tract. In mammals, cytokines and prostaglandins synthesized in male accessory glands induce changes in female gene expression that modify the structure of the endometrium to facilitate embryo development and implantation (Robertson, 2007).

Serotonin (5-HT) is an important modulator of female sexual function, in addition to its numerous other functions including mood regulation, behavioral inhibition, patience, reward, punishment, social behavior, defensive behavior, sensory gain, and learning (Soubrié, 1986; Deakin and Graeff, 1991; Clarke et al., 2004; Nakamura et al., 2008; Matias et al., 2017; Miyazaki et al., 2012; Miyazaki et al., 2014; Dugué et al., 2014; Fonseca et al., 2015; Cohen et al., 2015; Li et al., 2016; Seo et al., 2019). In human studies, more than 25% of women administered selective serotonin reuptake inhibitors (SSRIs) report sexual dysfunction, including decreased libido and difficulty achieving sexual arousal and orgasm (Lorenz et al., 2016). In rodents, SSRI administration suppresses behaviors such as lordosis that indicate sexual receptivity (Matuszczyk et al., 1998; Uphouse et al., 2006). In male mice 5-HT neural activity has been investigated during mating, and rises at mount onset (Li et al., 2016), but mating-related 5-HT neural activity in females has not been described. Neural activity in female mice that is associated with male ejaculation is of particular interest, because in mice ejaculation is required for the induction of pregnancy and pseudopregnancy (Yang et al., 2009).

Here, we describe DRN 5-HT neural dynamics in female mice during the consummatory phase of mating, which includes mounting, intromission (penile insertion), ejaculation, and post-ejaculation freezing. We use fiber photometry to record DRN 5-HT population neural activity in female mice during the complete mating sequence and in response to artificial mechanical and chemical sexual stimulation. Our findings reveal a pronounced increase in female DRN 5-HT neural activity that peaks specifically upon male ejaculation and which persists at a level above baseline until the male disengages. Using newly-developed surgical interventions to perturb specific aspects of ejaculatory physiology in males, we show that mechanical intravaginal stimulation provided by distal penis expansion (“penile cupping”) at ejaculation or forceful fluid expulsion are each sufficient to support ejaculation-related DRN 5-HT neural activity in recipient females without chemosensory cues.

## RESULTS

### Female DRN 5-HT neural activity increases at ejaculation

The consummatory phase of mating can be subdivided into events defined by male behavior, and here we investigate four—mount, intromission, pre-ejaculatory shudder, and post-ejaculatory freeze (**Figure 1A**). Mount is defined here as contact between the male abdomen and the back of the female, intromission is intravaginal penis insertion, pre-ejaculatory shudder is high-frequency thrusting that ends with ejaculation (**Video S1)**, and post-ejaculatory freeze is the prolonged immobile male grasping of the female following ejaculation (McGill, 1962). We used fiber photometry (Gunaydin et al., 2014) to monitor DRN 5-HT population neural activity in freely behaving female mice during mating. We injected AAVdj-EF1α-DIO-GCaMP6m (Chen et al., 2013) into the DRN of female SERT-Cre mice (Zhuang et al., 2005) and implanted an optical fiber over the DRN (**Figure 1B**). To control the timing and extent of female sexual receptivity during behavioral testing, we ovariectomized females and used a standard estradiol-progesterone priming protocol prior to mating (**Figure 1B**).

**Figure 1.**
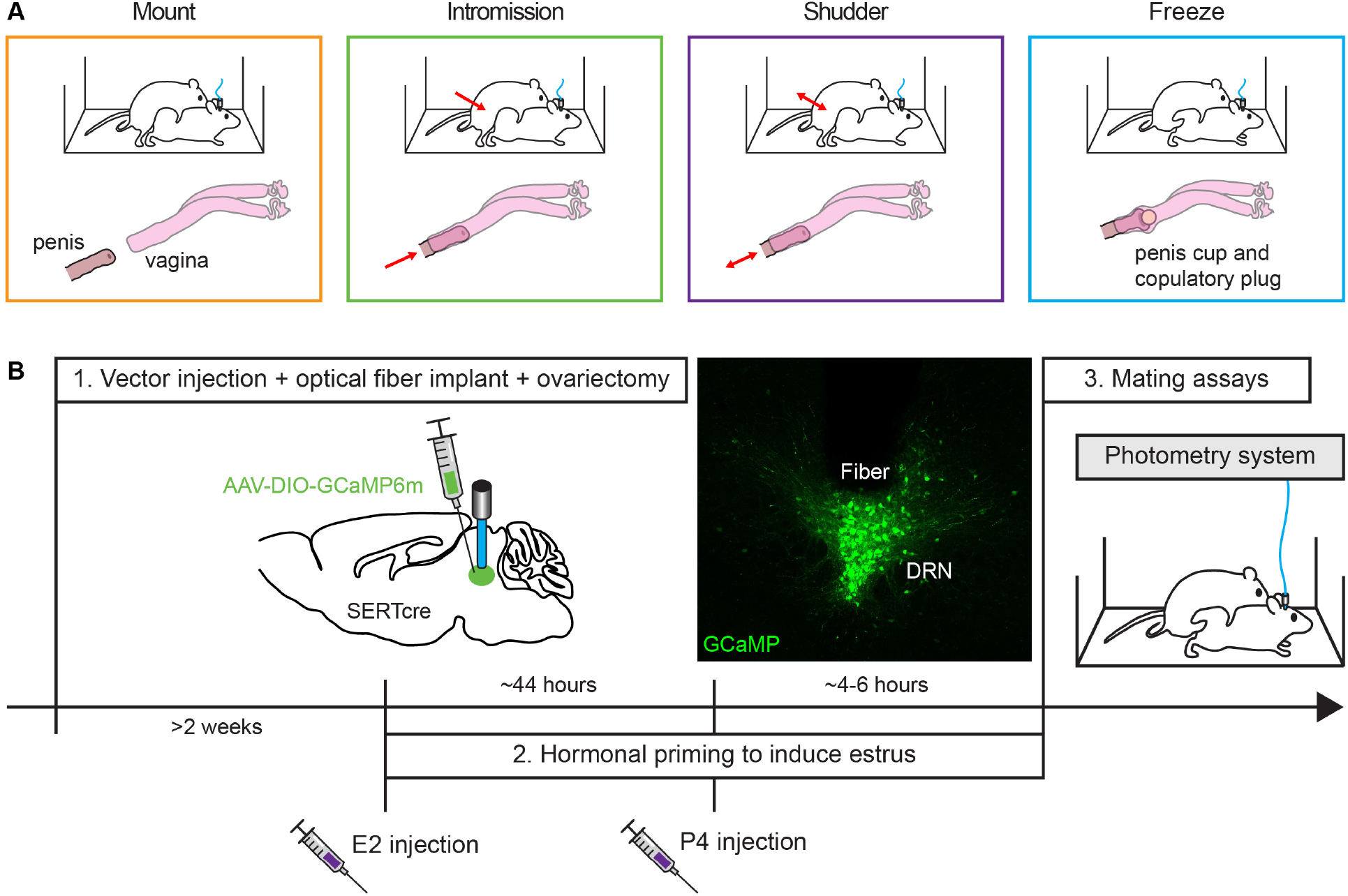
DRN 5-HT neural activity recorded with fiber photometry during mating. (A) Schematic of male sexual behaviors during the consummatory phase of mating. (B) Experimental timeline and GCaMP expression in DRN 5-HT neurons of a SERT-Cre female mouse. E2 = estradiol. P4 = progesterone.

Ejaculation was associated with a robust increase in female DRN 5-HT activity (GCaMP, n = 14 females, p < 0.001; GFP, n = 3 females, p = 0.75; Wilcoxon signed rank, **Figure 2 and Video S1**). This increase in neural activity peaked at ejaculation, and persisted above baseline for several seconds until a reduction was seen at the end of the post-ejaculatory intromission (GCaMP, n = 8 females, p < 0.05, paired t-test, D’Agostino-Pearson normally distributed, **Figures S1A and S1B**). Female DRN 5-HT activity did not significantly increase on pre-ejaculatory mounts or intromissions (mount, n = 7 females, p = 0.06; intromission, n = 7 females, p = 0.44; ejaculation, n = 8 females, p < 0.01; Wilcoxon signed rank test, **Figures S1C and S1D**). We also recorded DRN 5-HT neural activity in naturally cycling females during mating and found activation at ejaculation, indicating that this result did not depend on ovariectomy and hormonal priming (n = 4 females, p < 0.05, paired t-test, D’Agostino-Pearson normally distributed, **Figures S1E-H)**. DRN 5-HT activation upon ejaculation did not depend on female sexual experience (virgin, n = 9 females, p < 0.01; second mating, n = 5 females, p < 0.01; third mating, n = 5 females, p < 0.05; fourth mating, n = 7 females, p < 0.01, paired t-test, D’Agostino-Pearson normally distributed, **Figures S1I-K).** Thus, DRN 5-HT neural activity in female mice is specifically elicited by male ejaculation during the consummatory phase of mating.

**Figure 2.**
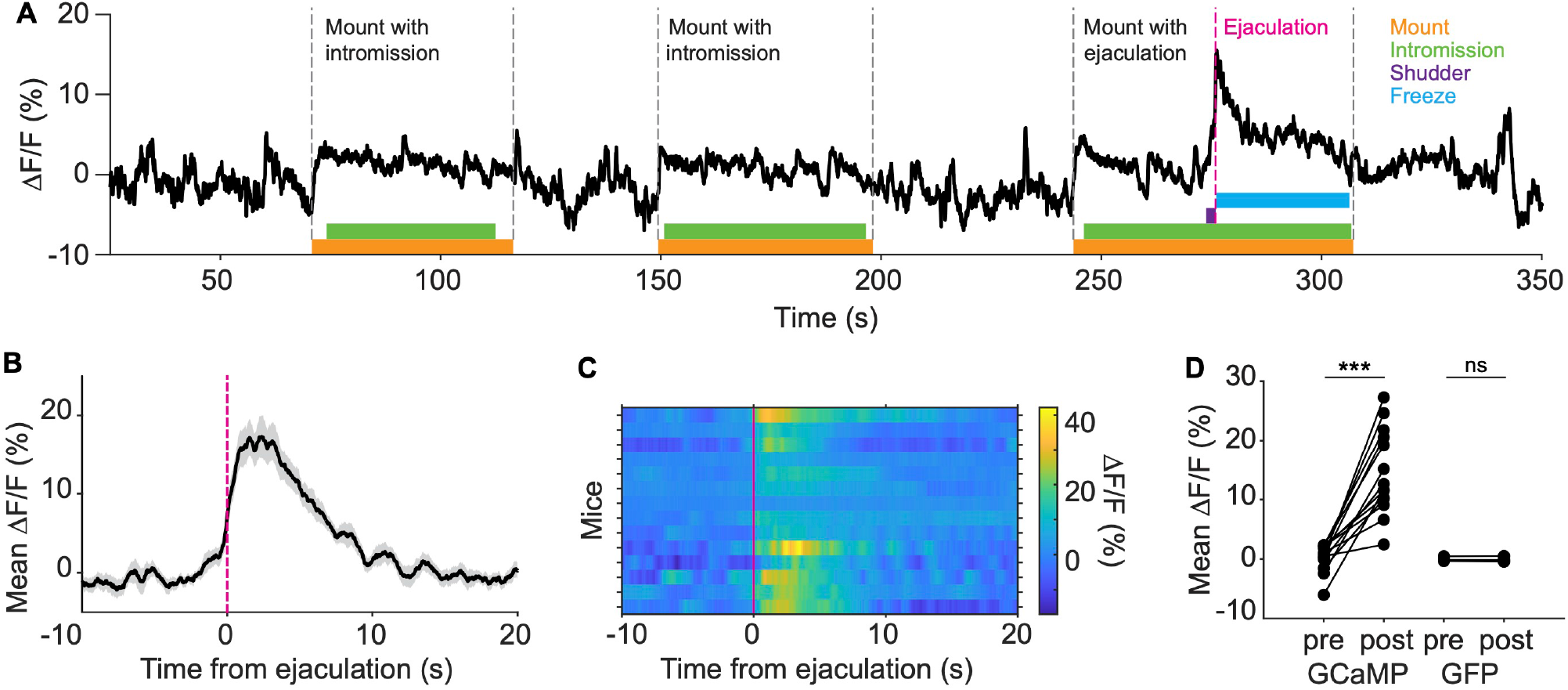
Female DRN 5-HT neural activity increases at ejaculation. (A) Example female DRN 5-HT GCaMP ΔF/F during mating. Colored boxes indicate scored male behaviors. Grey dotted lines indicate the onset and offset of male mounts. Magenta dotted line indicates ejaculation. (B) Mean DRN 5-HT GCaMP ΔF/F aligned to ejaculation (n = 14 females). Shaded area indicates SEM. (C) Heatmap of DRN 5-HT GCaMP ΔF/F for all animals, aligned to ejaculation (n = 14 females). (D) Mean DRN 5-HT ΔF/F 5 s before and after ejaculation in GCaMP (n = 14 females; mean diff = 14.25%) and GFP females (n = 3 females; mean diff = 0.03%) (Wilcoxon signed rank test, ns = non-significant, ***p < 0.001). See also Figure S1 and Video S1.

### Artificial mechanical intravaginal stimulation elicits increased DRN 5-HT neural activity

As ejaculation is required for the initiation of neuroendocrine processes in female mice that support pregnancy and pseudopregnancy (Yang et al., 2009), we sought to characterize the ejaculation-specific sensory stimuli that may be detected intravaginally by the female. Mechanical stimulation is provided by ejaculation-specific changes in the shape of the mouse penis and by the forceful expulsion of ejaculatory fluid or copulatory plug (McGill and Coughlin, 1970; Sachs, 1980). Additionally, sperm are accompanied by fluid that contains a variety of signaling molecules including prostaglandins and sex peptides (Rodríguez-Martínez et al., 2011; Avila et al., 2011), and chemical stimulation in some species is necessary for normal female reproductive behavior (Aprison and Ruvinsky, 2019; Chen et al., 1988; Yapici et al., 2008).

To investigate whether chemical or mechanical stimulation might underlie the increase in female DRN 5-HT neural activity upon ejaculation, we used a syringe to pseudo-randomly deliver 5 different stimuli into the vaginal canal of gently restrained female mice (**Figure 3A**). To test mechanical stimulation, we used a piece of rubber attached to the end of a syringe which could be inflated into a small balloon (McGill, 1970). To test chemical stimulation, we used two solutions: ejaculatory fluid (including the homogenized copulatory plug) collected from a female uterus immediately after ejaculation, and homogenized accessory sex glands from a male. As controls, we delivered equivalent volumes of air or saline. Fluid delivery was slow (0.1 mL at ∼0.2 mL/s), but may have provided a small amount of mechanical stimulation.

**Figure 3.**
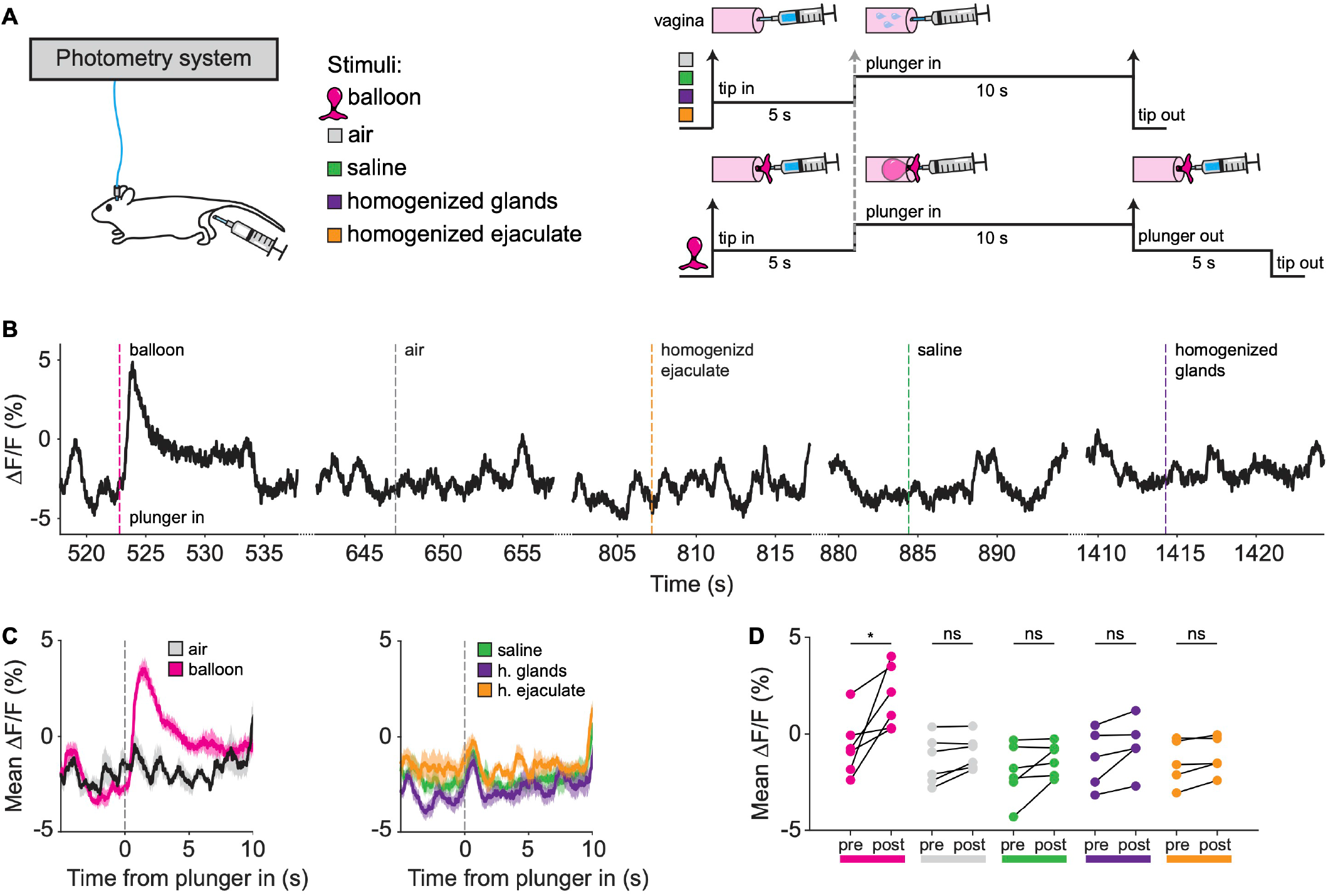
Artificial mechanical intravaginal stimulation elicits increased DRN 5-HT neural activity. (A) Experimental design. Artificial stimuli were delivered into the vagina of gently restrained females using a syringe. In each session each stimulus was delivered 5-10 times pseudo-randomly every >20 s. For all conditions, after stimulus delivery the syringe was held in place for ∼10 s. (B) Example DRN 5-HT GCaMP ΔF/F during artificial stimulation. Dotted lines for each condition indicate when the syringe plunger was pushed in. (C) Mean DRN 5-HT GCaMP ΔF/F aligned to plunger in for the same mouse for all conditions (n = 10 trials per condition). Shaded area indicates SEM. (D) Mean DRN 5-HT GCaMP ΔF/F 5 s before and after plunger in (balloon (mean diff = 2.51%), air (mean diff = 0.43), saline (mean diff = 0.67), n = 6 females; homogenized glands (mean diff = 0.69), homogenized ejaculate (mean diff = 0.32), n = 5 females) (Wilcoxon signed ranked test, ns = non-significant, *p < 0.05). See also Figure S2.

We observed a significant increase in DRN 5-HT neural activity in response to balloon inflation, but not in response to any of the other stimuli (GCaMP: balloon, n = 6 females, p < 0.05; air, n = 6 females, p = 0.09; saline, n = 6 females, p = 0.09; homogenized glands, n = 5 females, p = 0.06; homogenized ejaculate, n = 5 females, p = 0.13; Wilcoxon signed ranked test, **Figures 3B-3D**. GFP: balloon, n = 3 females, p = 0.59; air, n = 3 females, p = 0.82, paired t-test, D’Agostino-Pearson normally distributed, **Figures S2A and S2B**). DRN 5-HT neural activity rose at the moment the syringe plunger was depressed to inflate the balloon and fell when the plunger was pulled to deflate the balloon (balloon, n = 6 females, p < 0.05, paired t-test, D’Agostino-Pearson normally distributed, **Figures S2C and S2D**), supporting a strong association between DRN 5-HT neural activity and mechanical stretch caused by intravaginal balloon inflation.

There was a small but not significant rise in DRN 5-HT neural activity upon delivery of saline or liquid ejaculate stimuli (**Figure 2C**), which may be due to mechanical stimulation. Supporting this possibility, we observed that fast delivery of a larger volume of saline (0.7 mL at ∼0.7 mL/s) into the vaginal canal was sufficient to elicit female DRN 5-HT neural activity (**Figures S2E and S2F**). Hormonal priming did not influence the DRN 5-HT response to balloon inflation (not primed, n = 5 females, p < 0.05; primed, n = 5 females, p < 0.05, paired t-test, D’Agostino-Pearson normally distributed, **Figures S2G and S2H**). These data are consistent with the hypothesis that mechanical but not chemical intravaginal stimulation elicits activity in DRN 5-HT neurons in female mice.

### Separate blocking of penile cupping or ejaculatory fluid/copulatory plug expulsion does not disrupt female DRN 5-HT neural activity at ejaculation

Ejaculation is a dynamic process comprising several intravaginal steps. One step of potential relevance is the striking morphological change that occurs in the mouse penis at the moment of ejaculation (McGill and Coughlin, 1970). 2-3 seconds after the onset of the pre-ejaculatory shudder, the diameter of the distal end of the penis dramatically increases, forming a “penile cup” that lasts 5-12 seconds. This phenomenon has also been observed during *ex copula* induction of penile reflexes (Sachs, 1980). Following penile cupping, the ejaculatory fluid is forcefully expelled and a large copulatory plug is shot from the penis with such force that it embeds deeply into the cervix and occasionally penetrates the uterus (McGill and Coughlin, 1970). These ejaculation-specific mechanical stimuli are thought to induce vaginal/cervical stretching, and have been hypothesized to play a role in the induction of pseudopregnancy in mice (McGill and Coughlin, 1970). We sought to determine if these stimuli might contribute to the rise in female DRN 5-HT neural activity on male ejaculation.

We first investigated whether intravaginal mechanical stimulation during ejaculation might be provided by penile cupping (McGill and Coughlin, 1970). Transection of the paired bulbocavernosus (BC) muscles at the base of the penis has been shown to damage penis erectile function in mice, including *ex copula* penile cup display (Elmore and Sachs, 1988). To test whether penile cupping elicits the rise in female DRN 5-HT neural activity at ejaculation, we performed unilateral transections of the BC muscle in male mice (cup males, **Figure 4A**). Unilateral transections were selected over bilateral ones because we observed that male mice with complete transection of the BC muscles were unable to perform intromissions and ejaculations (data not shown). Males with unilateral BC transections displayed normal mating behavior, but did exhibit a greater number of “intromission breaks”—periods after intromission when the penis leaves the vagina and is reinserted without a dismount (Elmore and Sachs, 1988) (**Figure S3**). This behavioral phenotype results from damage to the penis glans erectile function, which is supported by the BC muscles and required for both continuous intromission and penile cup formation at ejaculation. We recorded DRN 5-HT neural activity from females mated with males with unilateral BC transections, and found a significant increase in neural activity at ejaculation, similar to the increase observed in the same females during mating sessions with unmodified control males (cup, n = 5 females, p < 0.05; control, n = 5 females, p < 0.05; paired t-test, D’Agostino-Pearson normally distributed, **Figures 4B and 4C**). Thus, disrupting penile cupping alone does not prevent the rise in female DRN 5-HT neural activity upon ejaculation.

**Figure 4.**
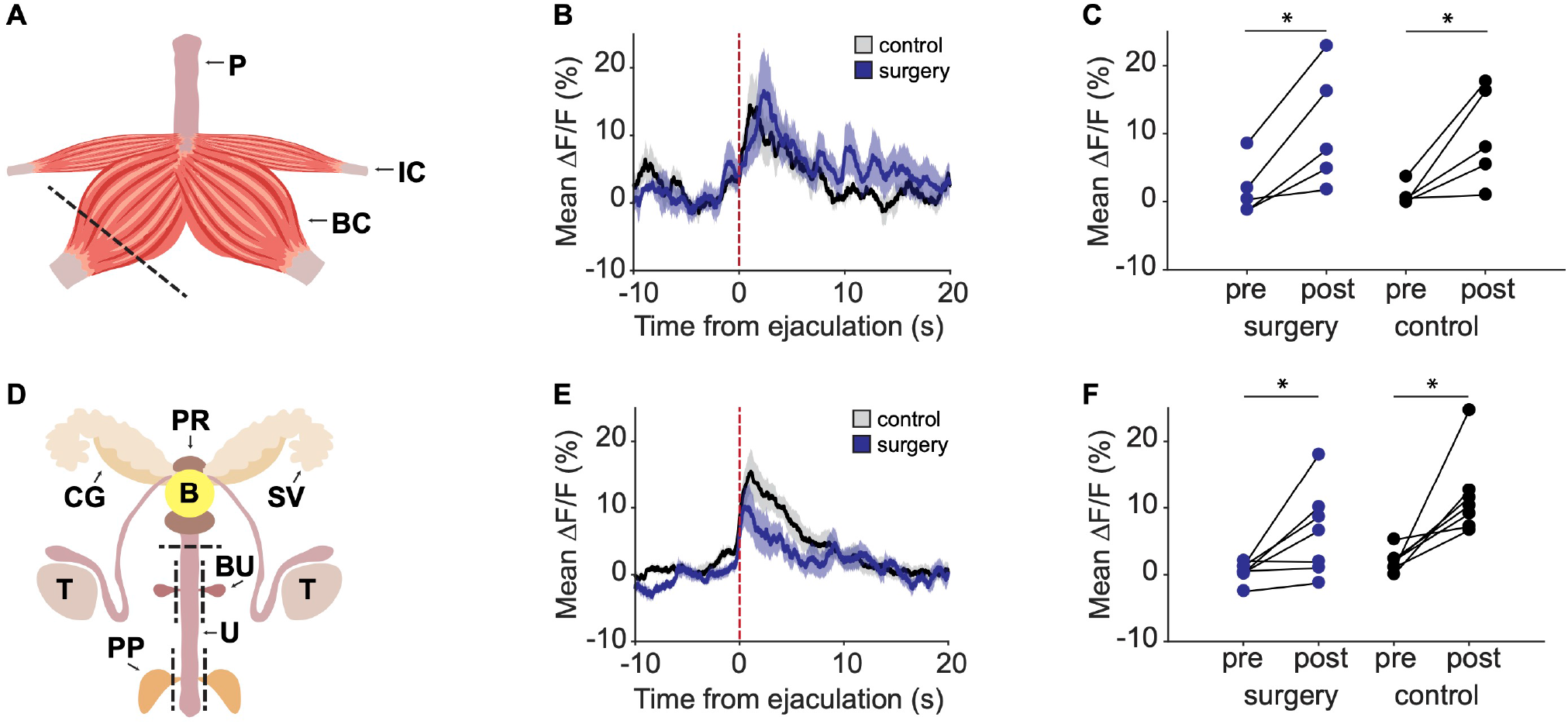
Separate blocking of penile cupping or ejaculatory fluid/copulatory plug expulsion does not disrupt female DRN 5-HT neural activity at ejaculation. (A) Surgical strategy to block penile cupping (cup males). The male right bulbocavernosus muscle located at the base of the penis was transected. BC = bulbocavernosus muscle, P = penis, IC = ischiocavernosus muscle. (B) Mean DRN 5-HT ΔF/F aligned to ejaculation in females (n = 5) mated with cup males (blue) and control males (black). Shaded area indicates SEM. (C) Mean DRN 5-HT ΔF/F 5 s before and after ejaculation in females (n = 5) mated with cup males (blue; mean diff = 9.01%) and control males (black; mean diff = 8.72%) (paired t-test, *p < 0.05; D’Agostino-Pearson normally distributed). (D) Surgical strategy to divert ejaculatory fluid/copulatory plug expulsion (fluid-plug males). B = bladder, T = testes, U = urethra, PR = prostate glands, BU = bulbourethral glands, PP = preputial glands, SV = seminal vesicles, CG = coagulating glands. (E) Mean DRN 5-HT ΔF/F aligned to ejaculation in females (n = 7) mated with fluid-plug males (blue) and control males (black). Shaded area indicates SEM. (F) Mean DRN 5-HT ΔF/F 5 s before and after ejaculation in females (n = 7) mated with fluid-plug males (blue; mean diff = 5.97%) and control males (black; mean diff = 9.80%) (Wilcoxon signed rank test, *p < 0.05). See also Figures S3 and S4.

Another possible source of mechanical stimulation is the forceful expulsion of the ejaculatory fluid and copulatory plug characterized in *ex copula* observations of the mouse ejaculatory sequence (McGill and Coughlin, 1970). To test whether ejaculatory fluid/copulatory plug expulsion elicits the rise in female DRN 5-HT neural activity at ejaculation, we surgically modified male mice to re-route urine and ejaculate through an alternative, non-penile exit (fluid-plug males, **Figure 4D and Video S2**). To generate these males, we transected the urethra caudal to the prostate glands and reconnected the cranial urethral segment to the abdominal wall, leaving a stoma through which urine and ejaculatory fluid could be diverted. Additionally, the bulbourethral and preputial glands—two pairs of accessory sex glands connected to the caudal urethral segment that also contribute components to the ejaculatory fluid—were resected. At the end of every mating session we verified that no copulatory plug was left inside the female vagina after shudder and freezing behavior. In a few instances, the copulatory plug was found on the female’s flank or perineal fur (**Figure S4**). We recorded DRN 5-HT neural activity from females mated with males with diverted ejaculatory fluid/copulatory plug, and found that there was significant DRN 5-HT neural activation at the time of ejaculation, similar to the signal observed in the same females during mating sessions with unmodified males (fluid-plug, n = 7 females, p < 0.05; control, n = 7 females, p < 0.05; Wilcoxon signed rank test, **Figures 4E and 4F**). Thus, disrupting ejaculatory fluid/copulatory plug expulsion alone does not prevent the rise in female DRN 5-HT neural activity upon ejaculation.

### Simultaneous blocking of both penile cupping and ejaculatory fluid/copulatory plug expulsion disrupts female DRN 5-HT neural activity at ejaculation

Because independently blocking the formation of the penile cup and the expulsion of the ejaculatory fluid/copulatory plug did not disrupt the rise in female DRN 5-HT neural activity upon ejaculation (**Figure 4**), we considered the possibility that blocking both factors simultaneously might be required. To test this hypothesis, we generated males with both surgeries—unilateral transection of the BC muscle to disrupt cupping, and urethral redirection to divert ejaculatory fluid/copulatory plug expulsion (cup/fluid-plug males, **Figure 5A**). When females mated with these males, we found no significant DRN 5-HT activation upon ejaculation (cup/fluid-plug, n = 7 females, p = 0.30; control, n = 7 females, p < 0.05; Wilcoxon signed rank test, **Figures 5B and 5C**). Male mating behavior was otherwise unchanged and indistinguishable from that of intact males mated with the same females, except for the expected display of intromission breaks (**Figure S3**). Thus, penile cupping or forceful expulsion of ejaculatory fluid each provide sufficient mechanical stimulation to elicit 5-HT neural activity during ejaculation, and simultaneous disruption of both was required to block this rise in neural activity.

**Figure 5.**
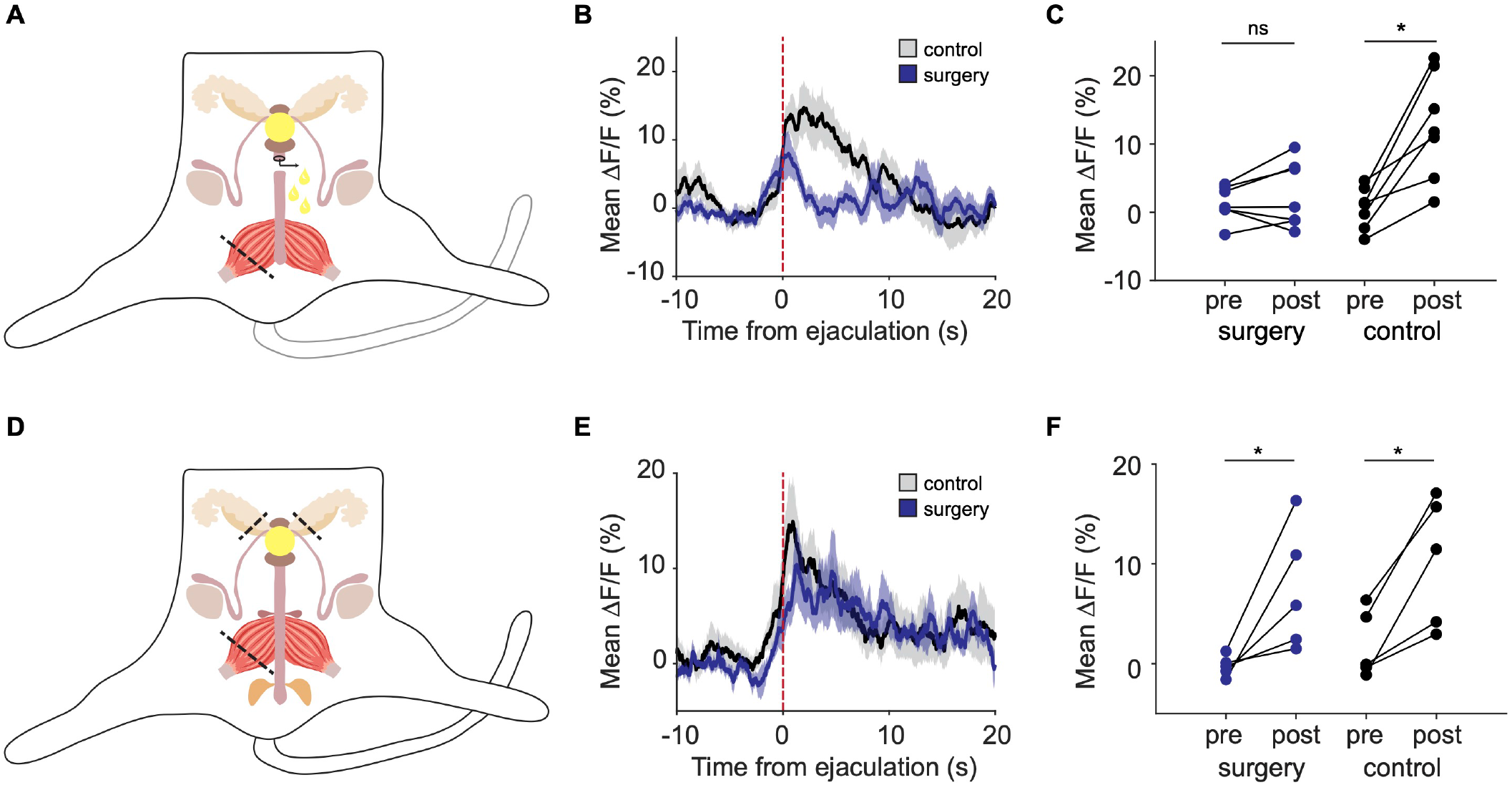
Simultaneous blocking of both penile cupping and ejaculatory fluid/copulatory plug expulsion disrupts female DRN 5-HT neural activity at ejaculation. (A) Surgical strategy to simultaneously block penile cupping and ejaculatory fluid/copulatory plug expulsion (cup/fluid-plug males). (B) Mean DRN 5-HT ΔF/F aligned to ejaculation in females (n = 7) mated with cup/fluid-plug males (blue) and control males (black). Shaded area indicates SEM. (C) Mean DRN 5-HT ΔF/F 5 s before and after ejaculation in females (n = 7) mated with cup/fluid-plug males (blue; mean diff = 1.27%) and control males (black; mean diff = 12.02%) (Wilcoxon signed rank test, ns = non-significant, *p < 0.05). (D) Surgical strategy to block penile cupping and copulatory plug expulsion, while preserving ejaculatory fluid expulsion (cup/plug males). (E) Mean DRN 5-HT ΔF/F aligned to ejaculation in females (n = 5) mated with cup/plug males (blue) and control males (black). Shaded area indicates SEM. (F) Mean DRN 5-HT ΔF/F 5 s before and after ejaculation in females (n = 5) mated with cup/plug males (blue; mean diff = 7.66) and control males (black; mean diff = 8.38%) (paired t-test, *p < 0.05; D’Agostino-Pearson normally distributed).

We observed a small but not statistically significant rise in female DRN 5-HT activity during ejaculation when females were mated with males carrying this combination surgery. One possible explanation for this trend is that female DRN 5-HT neurons may respond to tactile stimuli during ejaculation, such as the male postural change. Another possibility is that remnant penis erectile function was supported by the left BC muscle. As combination surgery males were able to perform intromissions, residual penile erectile function may have provided some mechanical stimulation at ejaculation. Consistent with this possibility, we also observed a statistically insignificant trend in the DRN 5-HT signal upon artificial delivery of fluid and air (**Figures 3C and 3D**). Future work could clarify the contribution of residual erectile function by measuring female DRN 5-HT responses during mating encounters with males carrying various degrees of muscle damage.

One question that emerges from our findings is how the expulsion of the ejaculatory fluid exerts mechanical force on the walls of the female reproductive tract. A feature of mouse ejaculatory fluid is its coagulation into a copulatory plug (Schneider, 2016), thus we sought to test whether the copulatory plug was necessary for female DRN 5-HT neural activation at ejaculation by interfering with coagulation. To block copulatory plug formation, we resected the coagulating glands and seminal vesicles, which produce a critical ingredient for copulatory plug formation, seminal vesicle secretion 2 (Gotterer et al., 1955; Kawano, 2014). We combined this with unilateral transection of the BC muscle to additionally block penile cupping (cup/plug males, **Figure 5D**). We verified at the end of every mating session that no copulatory plug was left in the female vagina. Females mated with these modified males exhibited robust DRN 5-HT neural activation at ejaculation (cup/plug, n = 5 females, p < 0.05; control, n = 5 females, p < 0.05; paired t-test, D’Agostino-Pearson normally distributed, **Figures 5E and 5F**), indicating that forceful ejaculatory fluid expulsion alone, without copulatory plug formation or penile cupping, provides enough mechanical stimulation to elicit female DRN 5-HT activity upon male ejaculation. Consistent with this finding, fast delivery of saline into the vaginal canal was sufficient to elicit female DRN 5-HT neural activity (**Figures S2E and S2F**). Together, these data reveal that mechanical stimulation provided either by male penile cupping or forceful fluid expulsion are sufficient to elicit an increase in female DRN 5-HT neural activity upon ejaculation.

## DISCUSSION

Here, we sought to investigate how female DRN 5-HT neurons respond to sexual stimulation during the consummatory phase of mating. Fiber photometry recordings from freely behaving females revealed pronounced DRN 5-HT neural activation elicited specifically by male ejaculation. Artificial intravaginal mechanical stimulation was sufficient to activate DRN 5-HT neurons, but intravaginal delivery of ejaculatory fluids was not. To identify the source of mechanical intravaginal stimulation responsible for female DRN 5-HT neural activation during natural mating, we developed surgical techniques to perturb two principal aspects of male ejaculatory physiology: forceful ejaculatory fluid expulsion and distal penis expansion (cupping) upon ejaculation. We found that both factors were independently capable of eliciting activity in female DRN 5-HT neurons, and that combined disruption of both was required to prevent the rise in female DRN 5-HT neural activity upon ejaculation.

### Mechanisms of vaginocervical stimulus detection

How does the female brain recognize sexual stimulation during mating and induce post-mating changes in female physiology and behavior? Uterine and vaginal sensation is primarily conveyed to the central nervous system via the pelvic and hypogastric nerves (Berkley et al., 1993; Hubscher and Berkley, 1995; Peters et al., 1987; Steinman et al., 1992). The pelvic nerve responds to stimulation of the vagina and cervix, while the hypogastric nerve responds to stimulation of the cervix and uterus (Peters et al., 1987; Berkley et al., 1990; Berkley et al., 1993). Pelvic nerve mechanosensory receptive fields are densely clustered at the fornix (the rostral end of the vaginal canal near the cervix), respond tonically to static pressure, and are activated by intravaginal balloon inflation (Berkley et al., 1990), while hypogastric nerve mechanosensory receptive fields are located in the cervix and the caudal body of the uterus and respond only to noxious mechanical stimulation intense enough to induce temporary ischemia (Berkley et al., 1988). Pelvic nerve function has been linked to sexual function while hypogastric nerve function plays an important role in childbirth and pelvic pain (Berkley et al., 1990).

Transection of the pelvic nerve blocks lordosis in response to vaginocervical stimulation (Carlson et al., 1965), shortens the duration of reduced sexual receptivity following intromission (Erskine, 1992), and prevents the induction of pseudopregnancy by vaginocervical stimulation (Kollar, 1953). However, combined transection of both pelvic and hypogastric nerves is required to block vaginocervical stimulation-induced immobility (Cunningham et al., 1991). The vagus nerve, which projects to the uterus (Ortega-Villalobos et al., 1990), may also play a role in conveying vaginocervical sensory information to the brain, as pupil dilation induced by vaginocervical stimulation is reduced but not abolished after spinal cord transection (Komisaruk et al., 1996).

Vaginocervical stimulation elicits changes in neural activity in a number of brain regions that have been implicated in the regulation of reproductive behavior and physiology. In female rats, mating with intromission induces cFos expression in a number of brain regions, including the preoptic area (POA), medial amygdala (MeA), and bed nucleus of the stria terminalis (BNST), and pelvic nerve transection blocks intromission-induced cFos induction in the POA and MeA (Rowe and Erskine, 1993). In female mice, mating stimulation sufficient to induce pseudopregnancy elicits cFos expression in the posterodorsal medial amygdala (MeApd), the ventrolateral part of the ventromedial hypothalamus (VMHvl), and the bed nucleus of the stria terminalis (BNST) (Yang et al., 2009). Progesterone receptor-expressing VMHvl neurons in female mice exhibit phasic responses to mounts and to ejaculation (Inoue et al., 2019), and cholecystokinin A receptor-expressing VHMvl neurons in female mice are persistently suppressed immediately following male ejaculation (Yin et al., 2022). In addition, BNST GABAergic neurons in female mice exhibit phasic responses specific to ejaculation (Bayless et al., 2019), and hypothalamic gonadotropin-releasing hormone (GnRH) neurons become persistently active following ejaculation in female mice (Wu et al., 1992). Neuromodulatory systems other than serotonin, such as dopamine and norepinephrine, have also been shown to respond to ejaculation in female mice (Etgen and Morales, 2002; Cameron et al., 2004; Dai et al., 2022). Investigation of the role of projection-specific DRN 5-HT neural activity in modulating ejaculation-related neural activity in these brain regions would be informative.

### DRN 5-HT neural activity and female sexual behavior

The DRN 5-HT system modulates a wide spectrum of functions—5-HT has been associated with behavioral inhibition, patience, reward, punishment, social behavior, defensive behavior, mood, sensory gain, and learning, among many others (Soubrié, 1986; Deakin and Graeff, 1991; Clarke et al., 2004; Nakamura et al., 2008; Matias et al., 2017; Miyazaki et al., 2012; Dölen et al., 2013; Miyazaki et al., 2014; Dugué et al., 2014; Fonseca et al., 2015; Cohen et al., 2015; Li et al., 2016; Seo et al., 2019). We have established a tight correspondence between female DRN 5-HT neural activity and male ejaculation, but we have not established a functional role for this rise in activity. Here, we consider several possibilities.

During ejaculation and the post-ejaculatory freeze, when DRN 5-HT neurons are active, female rodents maintain a stationary receptive posture that facilitates appropriate transfer of sperm and placement of the copulatory plug (Adler and Toner Jr, 1986; Dean, 2013; Elmore and Sachs, 1988; Matthews and Adler, 1977). 5-HT neural activity has been linked to behavioral inhibition in many but not all contexts (Soubrié, 1986, Seo et al., 2019), and the DRN 5-HT neural activation characterized here could play a role in facilitating immobility in response to sexual stimulation. The locomotor pause and receptive posture seen at ejaculation are also observed during other phases of mating that are not associated with significant DRN 5-HT neural activation, however, such as pre-ejaculatory intromissions, suggesting that DRN 5-HT neural activity in mating female mice plays a role specific to ejaculation and may function as a marker of potentially successful reproduction. Phasic 5-HT neural activity has also been linked to behavioral responses to emotionally-salient events, such as freezing and flight (Tessier et al., 2015; Seo et al., 2019; Paquelet et al., 2022), which may be relevant.

Alternatively, 5-HT may play a role in regulating the post-mating reduction in sexual receptivity in females, which is greater after ejaculation than after intromission without ejaculation (Peirce and Nuttall, 1961; Bermant, 1961; Bermant and Westbrook, 1966; Hill and Thomas, 1973; Erskine, 1985; Yang and Clemens, 1998). Decreased 5-HT function has been linked with a reduction in sexual receptivity—reducing 5-HT synthesis substantially impairs sexual receptivity in female flies (Ma et al., 2022), and female mice that lack 5-HT neurons prefer female odors over male odors (Zhang et al., 2013). Although we have not investigated this question here, long-term changes in DRN 5-HT neural activity following ejaculation, post-ejaculation desensitization of 5-HT receptors, or long-term state changes of DRN-5-HT-recipient circuits may play a role in this reduction in sexual receptivity.

5-HT and DRN neural activity has been linked to both reward and aversion (Simon et al., 1976; Deakin and Graeff, 1991; Nakamura et al., 2008; Dayan and Huys, 2008; Dayan and Huys, 2009; Dölen et al., 2013; McDevitt et al., 2014; Marcinkiewcz et al., 2016; Li et al., 2016; Seo et al., 2019, Paquelet et al., 2022), and may provide an emotional valence signal during mating. Mating is interesting, because while it is reinforcing—vaginocervical stimulation supports conditioned place preference (Meerts and Clark, 2009)—the post-mating reduction in immediate sexual receptivity suggests possible aversion that increases with increasing vaginocervical stimulation (McClintock et al., 1982). However, females could delay their return to the male for underlying reasons independent of the perceived valence of the mating event, such as stabilization of the copulatory plug (Adler and Toner Jr, 1986; McClintock et al., 1982). Much like the DRN 5-HT responses observed here, the pelvic nerve, which densely innervates the vaginal fornix, responds to gentle mechanical stimulation of the vagina and cervix associated with mating in rodents, including non-noxious balloon inflation (Berkley et al., 1990). However, the hypogastric nerve, which innervates the uterus and cervix, responds only to noxious mechanical stimulation associated with childbirth (Berkley et al., 1990; Berkley and Wood, 1995), and pelvic pain is treated with hypogastric nerve block, not pelvic (Plancarte et al., 1990). These findings suggest that the mechanical stimuli received by female mice upon male ejaculation may not be perceived as aversive by females.

Finally, we speculate that a potential role for ejaculation-related DRN 5-HT neural activity may be to enhance social olfactory learning and recognition in mice. During early pregnancy, female mice spontaneously abort if exposed to the odor of unfamiliar but not familiar males (Bruce, 1959), thus learning to recognize the odor of the mating male is of critical biological importance. DRN 5-HT neurons richly innervate the olfactory bulb (McLean and Shipley, 1987), and 5-HT has been implicated in odor processing (Petzold et al., 2009) and olfactory learning (McLean et al., 1993). Additionally, 5-HT contributes to the maintenance of early pregnancy in mice (Sahu and Dominic, 1980).

Overall, our findings support the hypothesis that a key regulator of DRN 5-HT activity is intravaginal mechanical stimulation provided by ejaculatory fluid expulsion and distal penis expansion. The surgical techniques developed here provide novel methods for the study of the functional impact of distinct aspects of ejaculation receipt on female physiology and behavior. Our results highlight the importance of exploring sex-specific functions of 5-HT for deepening our understanding of this neuromodulatory system.

## ACKNOWLEDGEMENTS

We thank J.R. Fetcho, J.H. Goldberg, N. Yapici, D. Lin, R.M. Harris-Warrick, I.T. Ellwood, R.J. Post, Y.Y. Ho, C.C. Vogt, C.H. Miller, W. Gu, D.A. Bulkin, B.J. Sleezer, and Y. Baumel for helpful discussions and/or comments on the manuscript; A.K. Recknagel for expert technical assistance; and Cornell Neurobiology and Behavior for training and support. Supported by the Mong Family Foundation (C.S., A.G.), the New York Stem Cell Foundation (M.R.W.), NIH DP2MH109982 (M.R.W.), the Alfred P. Sloan Foundation (M.R.W.), the Whitehall Foundation (M.R.W.), the Brain and Behavior Research Foundation (M.R.W.), and Cornell University.

## AUTHOR CONTRIBUTIONS

Conceptualization, E.L.T. and M.R.W.; Methodology, E.L.T. and M.R.W.; Software, C.S., A.G., and E.L.T.; Investigation, E.L.T. and M.R.W.; Analysis, E.L.T., C.S., A.G., and M.R.W.; Writing – Original Draft, E.L.T. and M.R.W.; Writing – Review and Editing, E.L.T., C.S., A.G., and M.R.W.; Supervision, M.R.W.

## STAR METHODS

## KEY RESOURCES TABLE

| REAGENT OR RESOURCE | SOURCE | IDENTIFIER |
| --- | --- | --- |
| **Virus Strains** |  |  |
| AAV-DJ-EF1α-DIO-GCaMP6m | Stanford Vector Core | N/A |
| AAV5-CAG-Flex-GFP | UNC Vector Core | N/A |
| **Chemicals** |  |  |
| Estradiol | Sigma-Aldrich | Cat # E8875; CAS 50-28-2 |
| Progesterone | Sigma-Aldrich | Cat # P0130; CAS 57-83-0 |
| DABCO | Sigma-Aldrich | Cat # 290734; CAS 280-57-9 |
| **Experimental Models: Organisms/Strains** |  |  |
| Mouse: C57BL/6J | The Jackson Laboratory | RRID:IMSR_JAX:000664 |
| Mouse: B6.129(Cg)-Slc6a4<sup>tm1(cre)Xz</sup>/J | The Jackson Laboratory | RRID:IMSR_JAX:014554 |
| **Software and Algorithms** |  |  |
| MATLAB | MathWorks | https://www.mathworks.com/products.html; RRID:SCR_001622 |
| GraphPad Prism | GraphPad Software | https://www.graphpad.com/scientific-software/prism/; RRID:SCR_002798 |

## RESOURCE AVAILABILITY

### Lead Contact

Further information and requests for resources and reagents may be directed to the Lead Contact, Melissa R. Warden.

### Materials Availability

This study did not generate new reagents.

### Data and Code Availability

Data and source code supporting the current study will be made available upon request.

## EXPERIMENTAL MODEL AND SUBJECT DETAILS

### Animals

All procedures conformed to guidelines established by the National Institutes of Health and have been approved by the Cornell University Institutional Animal Care and Use Committee. SERT-Cre female mice (The Jackson Laboratory, Bar Harbor, ME) between 2 and 12 months of age were used for 5-HT-specific viral vector expression. All Cre driver lines were fully backcrossed to C57BL/6J mice. All mice were maintained on a 12-hour reverse light-dark cycle with *ad libitum* access to food and water, and experiments were conducted during the dark portion of the cycle.

## METHOD DETAILS

### Viral vector injection and fiber implantation

Mice were anesthetized using isoflurane (5%). Fur was shaved and mice were placed in a stereotaxic frame (Kopf Instruments, Tujunga, CA). A heating pad was placed under the mice to prevent hypothermia. Isoflurane (1-2 %) was delivered via a nose cone throughout the surgery. Ophthalmic ointment was used to protect the eyes. Baytril (5 mg/kg, subcutaneous) and Ringer’s lactate solution (0.5 mL, subcutaneous) were given at the start of surgery. The scalp was disinfected with betadine scrub and 70% ethanol. A 1:1 mixture of 0.5 % lidocaine and 0.25% bupivacaine (0.1 mL) was injected intradermally along the incision line. An incision was made using a scalpel along the scalp midline. The exposed skull was thoroughly cleaned and a craniotomy was made above the DRN. The viral vector was targeted to the DRN (−4.5 AP, 0 ML, −3.3 & −2.9 DV), and pressure injected at 100 nl/min using a 10 µL Hamilton syringe (Nanofil; WPI, Sarasota, FL), 33 gauge beveled needle (Nanofil; WPI, Sarasota, FL), and micro-syringe pump controller (Micro 4; WPI, Sarasota, FL). After each injection, the needle was left in place for 10 minutes and then slowly withdrawn. A total of 800 nL (400 nL at each DV site) of viral vector was injected. A 400 µm diameter, 0.48 or 0.53 NA optical fiber (Doric Lenses, Quebec, Canada) embedded in a metal ferrule was implanted at 15° (−4.5 AP, 0.72 ML, −2.75 DV). A layer of metabond (Parkell, Inc., Edgewood, NY) and dental acrylic (Lang Dental Manufacturing, Wheeling, IL) was applied to firmly hold the implant in place. The surrounding skin was closed with a simple interrupted suture pattern. Subcutaneous buprenorphine (0.05 mg/kg), carprofen (5 mg/kg), and Ringer’s lactated solution (0.5 mL) were administrated post-operatively. Viral vector expression was allowed for a minimum of 2 weeks before behavioral testing.

### Ovariectomy

General surgical procedures were performed as described above. The ovariectomy protocol was conducted as reported previously (Souza et al., 2019), during the same surgical session as the viral vector injection and fiber implantation. Briefly, the dorsal fur between the last rib and the cranial pelvic edge was shaved on both sides. In each side, an incision was made into the peritoneal cavity, and the ovarian fat pad was carefully pulled out. The oviduct was clamped with hemostats, and the ovary was excised after cauterization of the tissue and blood vessels. The skin was sutured and covered with topical antibiotic (Water-Jel Technologies Llc., Carlstadt, NJ). Females were allowed to recover for at least 1 week before starting hormonal priming and behavioral testing.

### Urethrostomy

General surgical procedures were performed as described above. The ventral fur between the last rib and the penis base was shaved. An incision was made along the *linea alba* to access the peritoneal space. Using blunt dissection, the urethral segment caudal to the prostate gland was identified and pulled ventrally. The urethra was incised transversely, between the prostate and the dorsal penis curvature, taking care to not damage the pair of cavernous nerves running on both sides of the urethra. The caudal urethral end connected to the penis was cauterized and returned to the abdominal cavity. The sides of the cranial urethral wall connected to the bladder were sutured to the abdominal muscle and skin layers creating a stoma. The incision cranial and caudal to the stoma was closed, making sure that the urethral lumen remained continuous with the outside. Post-operatively, mice were kept on absorbent underpads to track normal urination for 3 days before being placed in normal bedding. Mice were allowed to recover for at least one week before behavioral testing. During this time, mice were checked daily, and common complications were monitored and treated appropriately, such as inflammation, infection, wound dehiscence, and urethral blockage. If animals presented signs of uroabdomen, they were euthanized immediately.

### Resection of accessory sex glands

General surgical procedures were performed as described above. The ventral fur between the last rib and the penis base was shaved, as well as the fur covering the scrotal sac. A midline incision was performed through the scrotal sac. Blunt dissection was used to identify the bulbourethral glands bilaterally. Each gland was pulled out, clamped at the base with a hemostat, and excised as close to its urethra connection as possible. The skin was sutured and covered with topical antibiotic.

Removal of the preputial glands was performed by making a midline incision into the skin layer ∼1 cm cranial to the penis base. After blunt dissection of the subcutaneous tissue, the preputial glands were pulled bilaterally. Each gland was clamped at the base with a hemostat, and excised. For removal of the seminal vesicles and associated coagulating glands, a midline incision along the *linea alba* was performed ∼1 cm cranial to the penis base. Blunt dissection was used to identify the seminal vesicles and coagulating glands, which were transected at their base connection to the bladder. All layers of the abdominal wall incision were sutured and covered with topical antibiotic.

### Partial bulbocavernosus (BC) muscle transection

General surgical procedures were performed as described above. The scrotal fur was shaved, and a midline incision was performed through the scrotal sac. Blunt dissection was used to identify the BC muscles and isolate them from the ischiocavernosus muscles. The right BC muscle was transected, and absorbable suture was placed in the cranial muscle end connected to the penis base. The scrotal skin incision was sutured and covered with topical antibiotic.

### Hormone priming

Estrus was induced in ovariectomized females with a subcutaneous injection of 10 µg β-estradiol 3-benzoate (E2) (Sigma) in 0.05 mL sesame oil ∼48 hours before testing. This was followed by a subcutaneous injection of 500 µg progesterone (P4) (Sigma) in 0.05 mL sesame oil 4 to 6 hours before testing. All females were ovariectomized and hormonally primed unless indicated otherwise.

### Staging of estrous cycle using vaginal cytology

Intact females were gently restrained to expose their vaginal opening. A 200 uL pipet tip connected to a rubber bulb was filled with ∼100 uL of saline solution kept at room temperature. The solution was flushed in and out of the vagina several times, making sure to not insert the pipet tip. Samples were placed on a microscope slide and allowed to air-dry. Samples were then stained using JorVet Dip Quick Stain (Jorgensen Labs, Loveland, CO) and examined under 10X magnification in a light microscope. The relative quantities of neutrophils, nucleated and anucleated epithelial cells were used to determine the estrous cycle stage (Cora et al., 2015). Female estrous cycle stage was scored for at least 2 weeks prior to behavioral testing during proestrus/estrus.

### Mating assay

Mating sessions were recorded at 30 frames/second under infrared illumination. Females were group housed and males were single housed for at least 1 week prior to testing. Both males and females had a range of sexual experience prior to the time of the behavioral assay. All mating sessions took place in the stud’s home cage, to which the female was transferred at the beginning of the session. The pair was allowed to interact for 15 min or until the stud performed the first successful intromission. If no intromission took place during this time, the pair was separated and the session was cancelled. After the first successful intromission, the female was connected to the fiber photometry system and recording started. Video and photometry data was recorded for up to 1 hour or until 3 minutes after successful ejaculation.

Videos were manually annotated using Bento (Segalin et al., 2020). Mounts were defined as the time when the female tail was between the male’s hindlegs and his abdomen was directly over the female dorsum. Intromissions scored from a side view video were defined as the time of low-frequency thrusting during mounts. Intromissions scored from bottom view videos were defined as the time when the penis was seen inside the vagina. Shudders were defined as the time of high-frequency thrusting that preceded freezing, where the male was completely immobile and connected to the female with all four limbs. Ejaculation was defined as the shudder offset. “Start to shudder on” latency was the time from the beginning of the photometry recording to the shudder onset. “Mount to shudder on” latency was the time from the onset of the last mount to the shudder onset. Failed mounts were defined as mounts with no intromission.

### Artificial vaginal stimulation

Six artificial stimuli were prepared and delivered 5-12 times each, in pseudorandom order. The combination of stimuli delivered in a given session varied by experiment. For all stimuli, a 1 mL tuberculin slip tip syringe connected to a ∼1.5 cm segment of PVC tubing (1/16” outer diameter) was inserted in the female vaginal canal. Care was taken to not exert pressure on the syringe beyond the natural resistance point of the caudal cervical wall. In the air condition, 0.1 mL of air were delivered. In the saline condition, 0.1 mL of saline were delivered at ∼0.2 mL/s. In the “forceful saline” condition, 0.7 mL of saline were delivered at ∼0.7 mL/s. For the gland fluid condition, all accessory sex glands and testes from a male mouse were collected, homogenized, and stored in saline solution at −80 degrees (Dean et al., 2009). In preparation for the artificial stimulation session, the solution was thawed to room temperature. For the uterus fluid condition, females were sacrificed immediately after mating, and their entire reproductive tract was extracted. The copulatory plug was extracted and homogenized in saline along with fluid stripped from the uterus. The solution was frozen at −80 degrees (Dean et al., 2011) and thawed to room temperature at the time of the artificial stimulation session. For the balloon condition, a ∼1×1 cm piece of balloon rubber was tied to the end of the tubing using suture material. The apparatus design was based on the one described in McGill, 1970. The syringe was filled with glycerol and labeled at the mark that resulted in inflation of a balloon with 6 mm diameter.

Females were gently restrained to expose their vaginal opening while artificial stimuli were delivered. Females were allowed at least 14 seconds of unrestrained rest in between deliveries, and at least 20 seconds in between stimuli. For all stimuli, the tip of the syringe tubing was inserted in the vaginal opening for ∼5 seconds before pressing the syringe plunger. In the empty, saline, gland fluid, and uterus fluid conditions, the syringe tubing was held inside the vagina for ∼10 seconds after pressing the plunger and before removal of the apparatus. In the balloon condition, the balloon was kept inflated inside the vagina for ∼10 seconds. After retracting the plunger, the syringe tubing with the deflated balloon was held in the vagina for ∼5 seconds before removing the apparatus. Artificial stimulation sessions were recorded at 30 frames/second under low lighting conditions, taking care to capture clear views of the female vaginal opening and the syringe plunger. Videos were manually annotated using Bento (Segalin et al., 2019).

### Perfusion and histological verification

Animals were deeply anesthetized with Fatal Plus at a dose of 90mg/kg, and transcardially perfused with 20 mL of phosphate-buffered saline (PBS), followed by 20mL of 4% paraformaldehyde (PFA) solution. Brains were extracted and stored overnight at 4°C in 4% PFA solution. After 24 hours, brains were transferred to 30% sucrose solution and allowed to equilibrate for a day. Brains were sectioned coronally (100 um) on a freezing microtome. Sections were washed in PBS and mounted on slides with PVA-DABCO. Images were acquired using a Zeiss LSM 800 confocal scanning laser microscope with a 20X air objective.

### Fiber photometry

The fiber photometry system used LEDs to deliver 405 nm and 473 nm light. These light streams were sinusoidally modulated at 211 Hz and 531 Hz respectively, combined into a minicube (Doric Lenses, Quebec, Canada), and coupled to an optical patch cord (400 µm; Doric Lenses). The optical patch cord was connected to the animal using a zirconia sleeve (2.5 mm; Doric Lenses). Emitted fluorescence was received through the same optical patch cord and collected by a femtowatt photodetector (New Focus 2151, Newport Corporation, Irvine, CA). This signal was processed using Doric Photometry Software. Photometry data analysis and statistical tests were performed using custom code in MATLAB (MathWorks, Natick, MA). Data were collected at 12 kHz, and low-pass filtered at 12 Hz. The calcium activity-independent 405 nm reference channel was fit to the calcium activity-dependent 473 nm channel using linear least squares. Relative fluorescence changes, reported as ΔF/F, were calculated using the following equation:

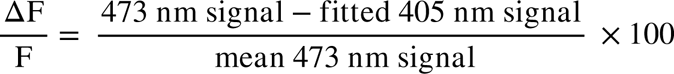

Mean ΔF/F before and after an event or behavior was calculated by calculating the mean signal in a 5 s window pre- and post-timepoint.

## QUANTIFICATION AND STATISTICAL ANALYSIS

### Statistics

Data are presented as mean ± standard error of mean (SEM), and statistical significance was considered for p < 0.05. Statistical analysis was performed using GraphPad PRISM (GraphPad Software Inc., CA). The magnitude of the change (mean diff) for each condition was calculated as the absolute value of the difference between the post and pre means. Sample distribution was assessed with a D’Agostino-Pearson omnibus normality test, and used to select parametric vs. non-parametric statistical tests for each dataset.

---

**Figure S1.**
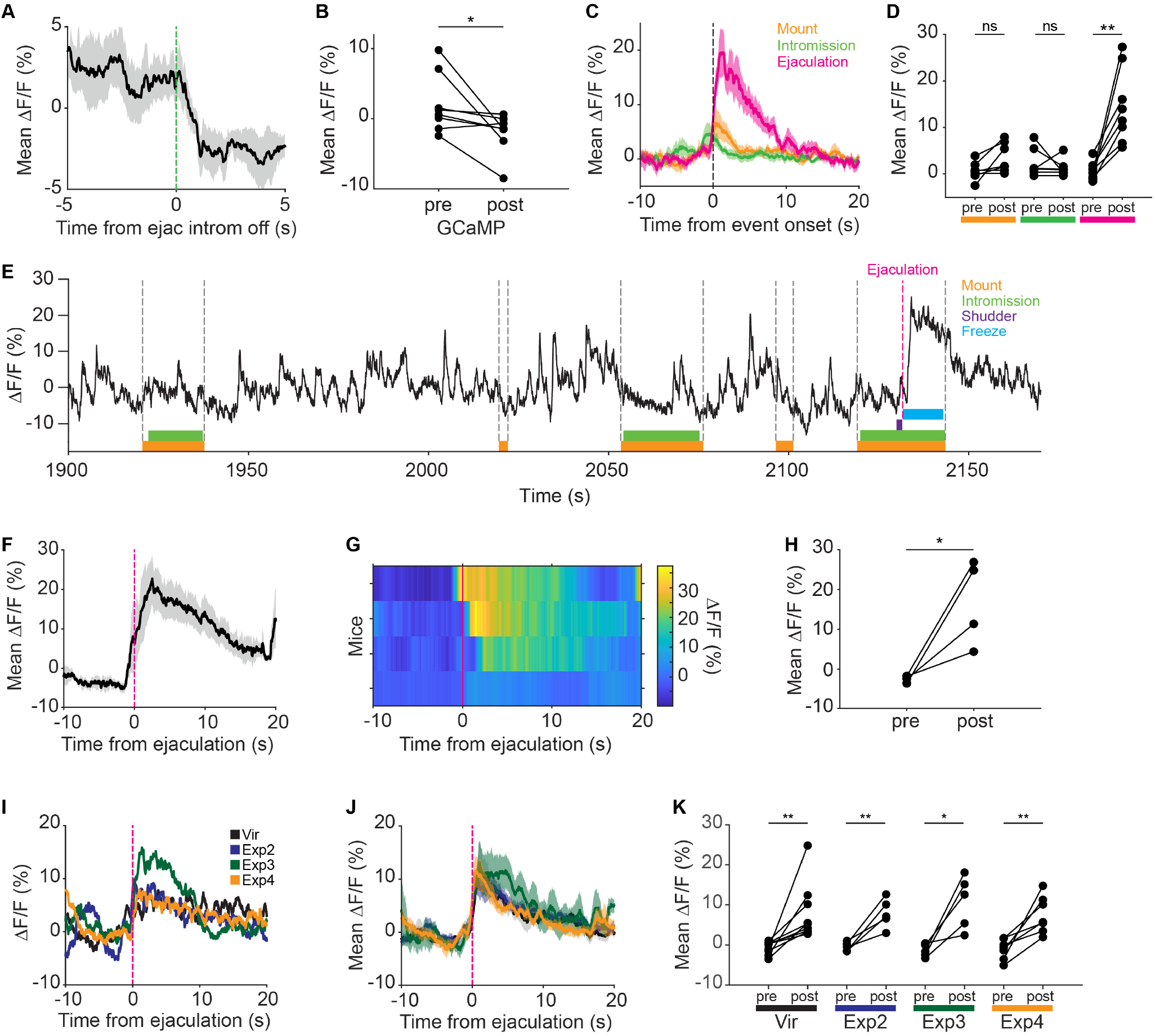
Female DRN 5-HT neural activity decreases at post-ejaculatory intromission offset, and DRN 5-HT neural activity is not elicited by pre-ejaculatory mounts and intromissions, related to Figure 2. (A) Mean DRN 5-HT ΔF/F aligned to post-ejaculatory intromission offset (n = 8 females). Shaded area indicates SEM. (B) Mean DRN 5-HT ΔF/F 5 s before and after male post-ejaculatory intromission offset (n = 8 females; mean diff = 4.08%) (paired t-test, *p < 0.05; D’Agostino-Pearson normally distributed). (C) Mean DRN 5-HT ΔF/F aligned to mount onset (orange, n = 7 females), intromission onset (green, n = 7 females), and ejaculation (magenta, n = 8 females). Shaded area indicates SEM. (D) Mean DRN 5-HT ΔF/F 5 s before and after mount onset (n = 7 females; mean diff = 2.83%), intromission onset (n = 7 females; mean diff = 1.12%), and ejaculation (n = 8 females; mean diff = 13.75%) (Wilcoxon signed ranked test, ns = non-significant, **p < 0.01). (E) Example DRN 5-HT ΔF/F for a naturally cycling female during mating. Color boxes indicate scored male behaviors. Grey dotted lines indicate the onset and offset of male mounts. Magenta dotted line indicates the moment of ejaculation. (F) Mean DRN 5-HT ΔF/F aligned to ejaculation for naturally cycling females (n = 4 females). Shaded area indicates SEM. (G) Heatmap of DRN 5-HT ΔF/F (n = 4 females) aligned to ejaculation. (H) Mean DRN 5-HT ΔF/F 5 s before and after ejaculation in naturally cycling females (n = 4 females; mean diff = 19.32%) (paired t-test, *p < 0.05; D’Agostino-Pearson normally distributed). (I) ΔF/F aligned to ejaculation for example ovariectomized female in her consecutive virgin (Vir, black), second (Exp2, blue), third (Exp3, green), and fourth (Exp4, orange) mating experiences. (J) Mean ΔF/F aligned to ejaculation for ovariectomized females in their virgin (black), second (blue), third (green), and fourth (orange) mating experiences. Shaded area indicates SEM. (K) Mean ΔF/F calculated in a 5 s window before and after ejaculation for ovariectomized females in their virgin (n = 9 females, **p < 0.01), second (n = 5 females, **p < 0.01), third (n = 5 females, *p < 0.05), and fourth (n = 7 females, **p < 0.01) mating experiences (paired t-test, D’Agostino-Pearson normally distributed).

**Figure S2.**
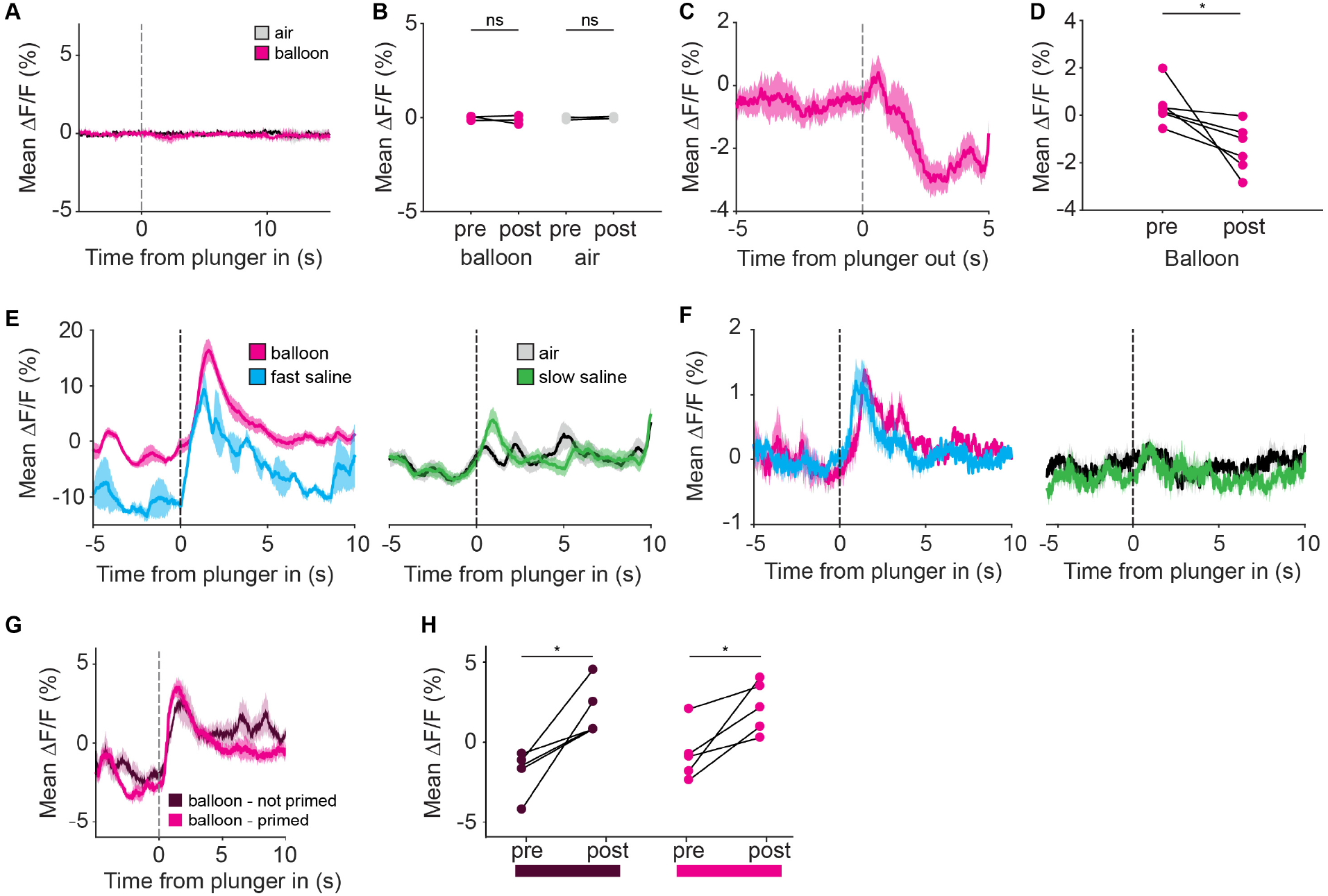
GFP ΔF/F during artificial mechanical intravaginal stimulation, and GCaMP ΔF/F during balloon deflation, related to Figure 3. (A) Mean DRN 5-HT GFP ΔF/F aligned to plunger in (n = 3 females). Shaded area indicates SEM. (B) Mean DRN 5-HT GFP ΔF/F 5 s before and after plunger in for balloon (n = 3 females; mean diff = 0.10%) and air (n = 3 females; mean diff = 0.01%) (paired t-test, ns = non-significant; D’Agostino-Pearson normally distributed). (C) Example mean DRN 5-HT GCaMP ΔF/F aligned to balloon plunger out (n = 10 trials). (D) Mean DRN 5-HT GCaMP ΔF/F 5 s before and after plunger out (n = 6 females; mean diff = 1.80%) (paired t-test, *p < 0.05; D’Agostino-Pearson normally distributed). Shaded area indicates SEM. (E) Mean DRN 5-HT ΔF/F aligned to plunger in for balloon inflation and fast saline in first example female, left. Air and slow saline, right. Shaded area indicates SEM. (F) Mean DRN 5-HT ΔF/F aligned to plunger in for balloon inflation and fast saline in second example female, left. Air and slow saline, right. Shaded area indicates SEM. (G) Mean DRN 5-HT GCaMP ΔF/F during artificial stimulation with balloon for an example female with and without hormonal priming. Dotted line indicates when the syringe plunger was pushed in. Shaded area indicates SEM. (H) Mean DRN 5-HT GCaMP ΔF/F calculated in a 5 s window before and after plunger in (balloon and not primed, n = 5 females; balloon and primed, n = 5 females; paired t-test, *p < 0.05; D’Agostino-Pearson normally distributed).

**Figure S3.**
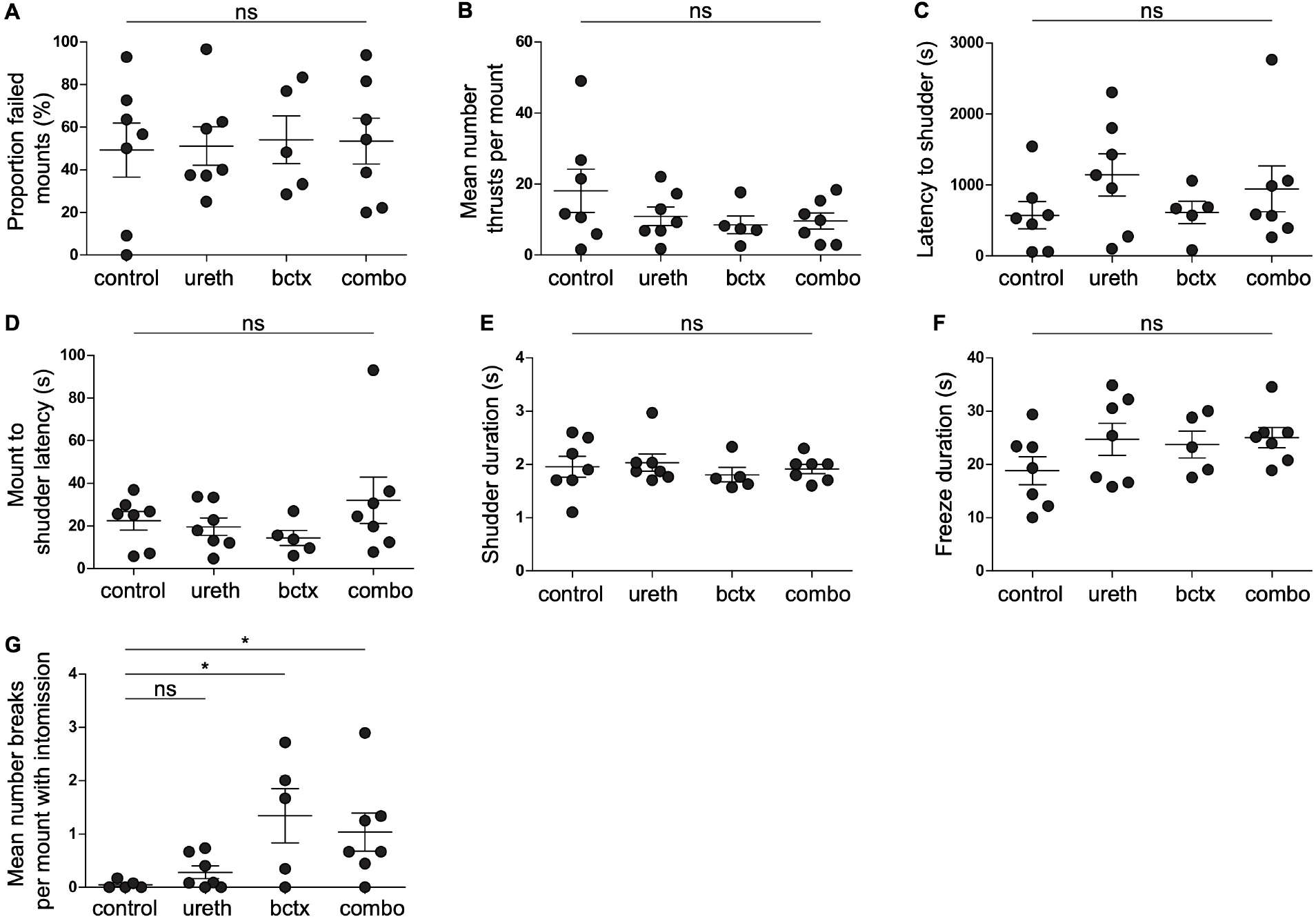
Surgically modified males exhibit normal mating behavior except for intromission breaks, related to Figures 4 and 5. (A) Percent mounts that did not result in intromission (one-way ANOVA, p = 0.99). Control (n = 7 males); ureth = urethrostomy to redirect ejaculatory fluids (n = 7 males); bctx = bulbocavernosus transection to disrupt penile cupping (n = 5 males); combo = both surgeries (n = 7 males). Each data point is a mating session. (B) Mean number of intromission thrusts per mount (one-way ANOVA, p = 0.32). (C) Time from start of session to shudder onset (one-way ANOVA, p = 0.39). (D) Time from start of final ejaculatory mount to shudder onset (one-way ANOVA, p = 0.37). (E) Duration of shudder (one-way ANOVA, p = 0.80). (F) Duration of post-ejaculatory freeze (one-way ANOVA, p = 0.29). (G) Mean number of intromission breaks per mount (one-way ANOVA, *p < 0.05). Only mounts with intromission were included.

**Figure S4.**
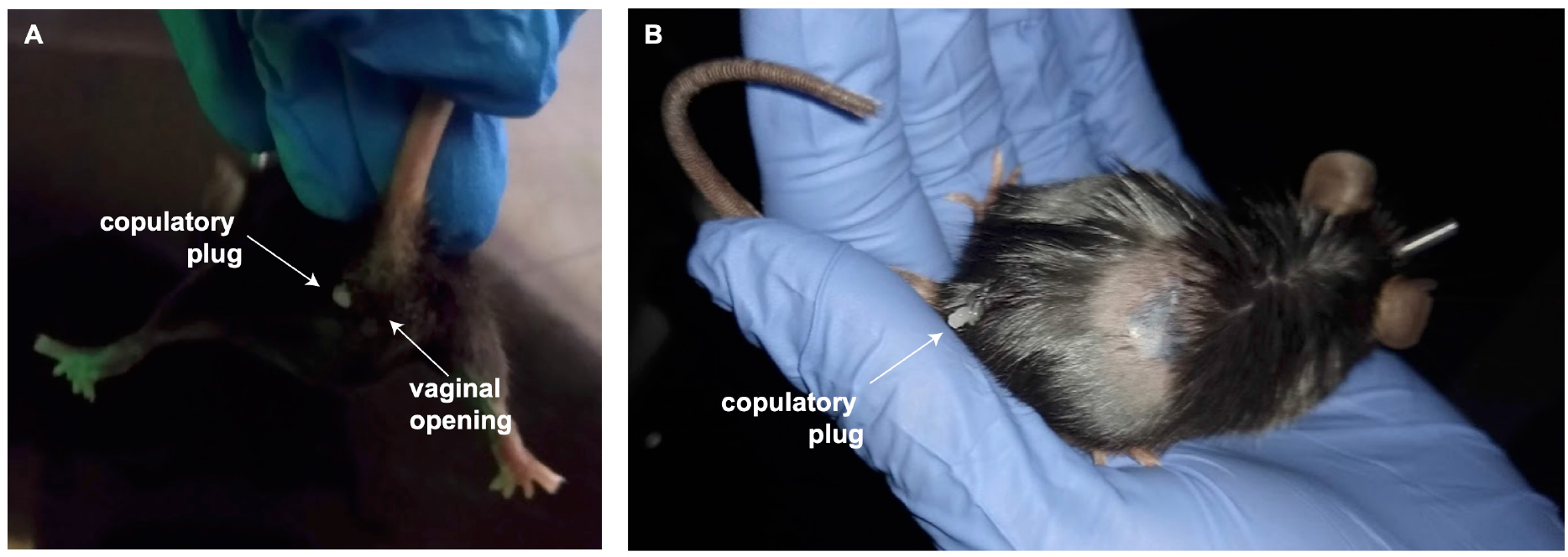
Copulatory plug found outside the vagina after mating sessions with urethrostomized males, related to Figure 4. (A) Copulatory plug found lateral to the vaginal opening in first example female immediately after mating session with urethrostomized male. (B) Copulatory plug found lateral to the tail base in second example female immediately after mating session with urethrostomized male.

**Video S1. Female DRN 5-HT neural activity increases at ejaculation, related to Figure 1.**

DRN 5-HT neurons in a SERT-Cre female show increase in GCaMP6 fluorescence at ejaculation, but not during pre-ejaculatory intromissions. Video frames are aligned to red line in female DRN 5-HT photometry trace. Color boxes indicate scored male behaviors.

**Video S2. Bladder expression in example urethrostomized male, related to Figure 4.**

Gentle pressure applied externally to the bladder of a urethrostomized male results in urine release via the artificial stoma instead of the penis.

